# Constitutive Siglec-1 expression by human dendritic cell precursors enables HIV-1 replication and transmission

**DOI:** 10.1101/578377

**Authors:** Nicolas Ruffin, Ester Gea-Mallorquí, Flavien Brouiller, Mabel Jouve, Aymeric Silvin, Peter See, Charles-Antoine Dutertre, Florent Ginhoux, Philippe Benaroch

## Abstract

Human dendritic cell (DC) lineage has been recently unraveled by high dimensional mapping revealing the existence of a discrete new population of blood circulating DC precursor (pre-DC also referred to as AS DC). Among all blood DC subsets, only pre-DC highly express Siglec-1, a lectin-like receptor able to bind HIV-1. We show that pre-DC are uniquely equipped among blood DC populations to promote HIV-1 replication and dissemination. Pre-DC stands out as the most susceptible DC population to infection by both HIV-1 CXCR4- and CCR5-tropic viral particles in a Siglec-1-dependent manner. HIV-1-infected pre-DC produce new viral progeny and transmit the virus to CD4^+^ T cells. Upon TLR activation, pre-DC become resistant to HIV-1 fusion and thus to infection and switch to a replication-independent mechanism of virus transfer to activated primary T lymphocytes mediated by Siglec-1. Thus, beside their role in DC ontogeny, blood pre-DC possess stage-specific properties that HIV-1 may exploit for viral spreading and modulation of the immune response.

## Introduction

Dendritic cell (DC) populations are heterogeneous and play a dual role with regards to HIV-1: key players in the establishment and spreading of HIV-1 infection^1,2^, DC also elicit innate and adaptive immune anti-viral responses^3–5^. Recently, development of high dimensional approaches has led to the identification of pre-DC, a new population of circulating precursors of DC^6,7^. The potential role of pre-DC in the course of HIV-1 infection remains to be established.

Dendritic cells (DC) are professional pathogen-sensing and antigen-presenting cells that are central to the initiation and regulation of immune responses^8^. The DC population is classified into two lineages: plasmacytoid DC (pDC), and conventional DC (cDC), the latter comprising cDC1 and cDC2 sub-populations^9^. Recently, DC ontogeny has been the object of intensive work using new high dimensional approaches including single cell level in mice^10^ and in humans^6,7,11^. Our study identified a new population of precursors of DC (pre-DC) that originates from bone marrow, circulate in the blood before reaching tissues where they can differentiate into cDC1 and cDC2^6^. Albeit human pre-DC share many markers with pDC, they are clearly different morphologically, phenotypically and functionally^6^.

DC are targets for HIV-1, although with lower efficiency than activated CD4^+^ T lymphocytes and macrophages^12,13^. DC are thought to participate in the establishment of the infection due to the expression of lectin-like receptors, allowing them to capture and transfer HIV-1 to CD4^+^ T cells^1,2^. Since the seminal study of Steinman, it is clear that DC possess the capacity to capture and transfer HIV particles with a high efficiency to T cells^12^. The process called trans-infection involves the lectin-like receptor Siglec-1 present on the surface of DC but still remains incompletely understood and essentially documented with *in vitro* derived dendritic cells from monocytes. Different mechanisms involving DCs in the early steps of the infection have been proposed but their precise role *in vivo* remains to be established.

The capacity of the various DC subsets to initiate anti-viral T-cell responses through antigen presentation and to respond to infection is highly organized in space and time^14^. pDC detect viruses, rapidly respond by producing large amounts of type I interferon (IFN), and may be, at least partially, responsible for the high levels of IFNα measured during the acute phase of HIV-1 infection^15^. cDC1 are resistant to HIV-1 infection, but can uptake HIV-1-infected cells to prime specific T-cell responses^14^. In contrast to pDC and cDC1, cDC2 are susceptible to HIV-1 infection *in vitro* and may thus provide, after dying and being phagocytosed, a source of viral antigens for cDC1 leading to cross presentation^14^.

With the recent discovery of the pre-DC subset, the question of its role as compared to the other blood DC subsets became of interest. Importantly, Siglec-1 (CD169) is one of the very few specific and constitutive markers of pre-DC (^7^ and Florent Ginhoux, personal communication). Siglec-1 is an interferon-induced receptor that can bind gangliosides, including GM3 that is part of the HIV-1 membrane and mediate efficient viral capture and transmission to T cells in vitro^16–20^. Hence pre-DC appears as a potentially critical population of cells with regards to HIV-1 infection prompting us to study its role as compared to the other blood DC subsets. We found that pre-DC are the most susceptible DC population to HIV-1 infection. Notably, both HIV-1 capture by pre-DC and HIV-1 infection of pre-DC are Siglec-1-dependent. Upon TLR-mediated activation, pre-DC up-regulate RAB15 expression and become resistant to HIV-1 fusion and thus to infection. Importantly, activated pre-DC switch to a replication-independent mechanism of virus transfer to activated primary T lymphocytes mediated by Siglec-1. We also demonstrate that, although being a minor fraction of peripheral DC, the depletion of pre-DC leads to an important decrease of HIV-1 infection in primary DC. These results show, for the first time, the potential involvement of pre-DC in HIV-1 dissemination to CD4^+^ T cells.

## Results

### Pre-DC is the most susceptible blood DC subsets to HIV-1 infection

We first set up an isolation strategy of the four blood DC subsets; pre-DC, pDC, cDC1 and cDC2, based on their differential surface protein expression, by a combination of magnetic bead negative selection and flow cytometry sorting (Figure S1a). The sorted populations had the expected^6,7^ frequency (Figures S1b and S1c) and morphology at the ultra-structural level (Figure S1D). Pre-DC (CD33^+^CD45RA^low^CD123^+^) represented roughly 1.5% of the total blood DC and our sorting procedure starting typically from 2.10^9^ PBMC allowed to obtain 10^5^ pre-DC (Figure S1c).

Among the specific markers of pre-DC, Axl and Siglec-1 stand out as the two most discriminative between DC subsets (Figures 1a and 1b,^6,7^). Given that Siglec-1 can bind HIV particles^16–18^, we evaluated pre-DC susceptibility to HIV-1 infection as compared to the other blood DC populations. For this purpose, these populations were purified from healthy donor blood according to our sorting strategy (Figure S1a) avoiding usage of the Siglec-1 marker given its potential involvement with HIV-1. When exposed to a CCR5-tropic HIV-1 encoding GFP (HIV-1 R5GFP), pre-DC and cDC2 were infected at a similar extent (mean±S.D. 4.2±2.5% and 3.2±2.6% of infected cells respectively after 48 h), while cCD1 and pDC remained refractory to infection (Figures 1c and 1d). Interestingly, a CXCR4-tropic HIV-1 (HIV-1 X4GFP) yielded higher rates of infection with pre-DC (4.9±1.4%) compared with any of the other subsets including cDC2, which all remained very low (Figure 1c and 1e). The low susceptibility of DC to HIV-1 infection has been linked to the presence of SAMHD1, a restriction factor that degrades cytosolic dNTPs, thus limiting the retro-transcription of the viral RNA and the infection rates^21–23^. However, no difference in SAMHD1 mRNA level was detected among all DC subsets, including pre-DC (Figure S2a). Due to the very low numbers of pre-DC isolated, we were unable to follow SAMHD1 phosphorylation that negatively regulate its activity^23,24^. Nevertheless, SAMHD1 activity can be efficiently counteracted by the accessory protein Vpx from HIV-2^21-23,25^. Accordingly, complementation of our HIV-1 strains with Vpx (using a Vpr-Vpx fusion construct) highly increased the rates of infection that reached up to 60% for pre-DC and approximately 20% for cDC2, both with HIV-1 R5GFP (Figures 1f and 1g), establishing that SAMHD1 remains in part active in these two populations, as observed with monocyte-derived macrophages (MDM)^21^. Importantly, with HIV-1 X4GFP complemented with Vpx, pre-DC was the only population infected (up to 49.5%, Figure 1f and 1h), further underlining the unique susceptibility of the pre-DC population to HIV-1 infection. Of note, monocyte-derived DC (MDDC) remained resistant to both viruses, and only slightly susceptible to HIV-1 R5GFP when complemented with Vpx (Figures S2b and S2c), as previously shown^12,26,27^.

**Figure 1.**
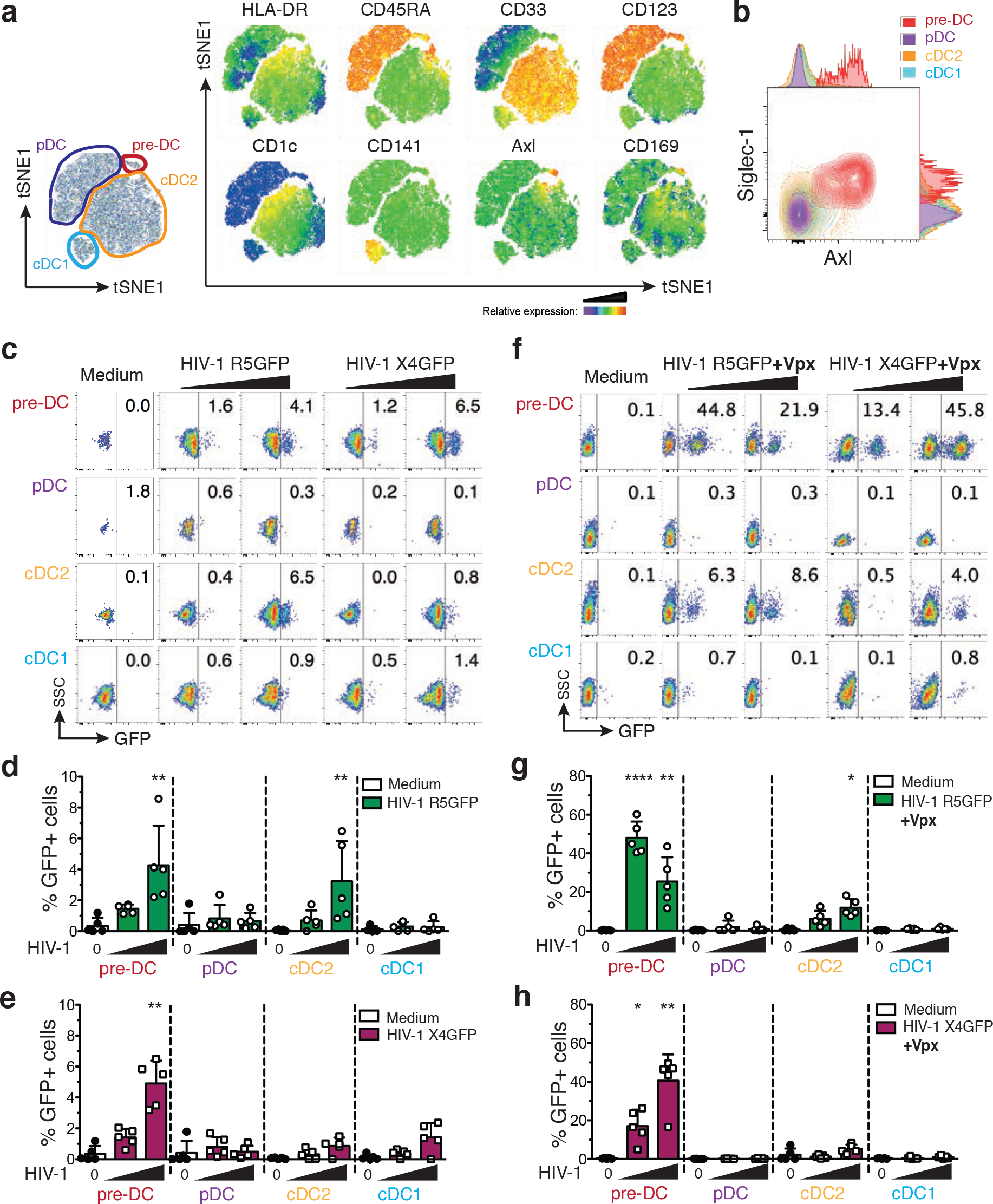
Comparative analysis of human blood DC subsets for their susceptibility to HIV-1 strains. **a**, Representative tSNE plots of flow cytometry data from HLA-DR^+^Lin(CD3/CD14/CD16/CD19/CD34)^−^ PBMC, showing gates defining the indicated DC subsets and relative expression of the indicated markers. **b,** Representative dot plot of Siglec-1 and Axl expression on pre-DC (HLA-DR^+^Lin^−^ CD33^int^CD45RA^+^CD123^+^Axl^+^), pDC (HLA-DR^+^Lin^−^CD33^-^CD45RA^+^CD123^+^Axl^-^), cDC2 (HLA-DR^+^Lin^−^CD33^+^CD45RA^-^CD123^-^CD1c^+^) and cDC1 (HLA-DR^+^Lin^−^ CD33^+^CD45RA^-^CD123^-^CD141^+^). **c,** GFP expression in blood DC subsets sorted and infected for 48 h with HIV-1 R5GFP or X4GFP. Freshly prepared viral preparations used for the experiments depicted from (c) to (h) were titrated on GHOST reporter cells in parallel and used at a MOI of 0.994 ± 0.112 (mean ± S.E.M). Representative dot plot analyses are presented with the percentages of GFP^+^ cells indicated in each dot plot. **d,** Quantification with HIV-1 R5GFP infection or **e,** with HIV-1 X4GFP as in (c), n=5 independent donors combined from 3 experiments. Individual donors are displayed with bars representing mean ± S.D. **f,** As in (c) using the indicated viruses complemented with Vpx. g, Quantification of infection with HIV-1 R5GFP or **h,** with HIV-1 X4GFP, both supplemented with Vpx, n=5 independent donors combined from 2 to 4 experiments. Individual donors are displayed with bars representing mean ± S.D..

We noticed that even in the presence of Vpx when using an R5-tropic HIV-1, cDC2 remained much less susceptible to the infection than pre-DC in the same conditions (up to 18 % versus 60 %, respectively (Figure 1g)), suggesting that besides SAMHD1, other mechanisms might limit the infection of cDC2.

### Specific Siglec-1 expression by pre-DC confers an advantage over other DC subsets for HIV-1 capture and infection

Since pre-DC represent only 1.5% of the total blood DC, the question of the contribution of this small population to the HIV-1 infection and spreading is of importance. To address this point *in vitro*, total DC were isolated from PBMC by enrichment using the pan-DC magnetic sorting procedure followed by flow cytometry sorting of all Lin^-^ HLA-DR^+^ cells that were used as “total DC”, or further sorted to obtain a DC population called “no pre-DC”, by excluding pre-DC based on their expression of CD123 and Axl. Both populations were exposed either to HIV-1 R5GFP or to HIV-1 X4GFP complemented with Vpx. The rates of infection, as measured at 48h post-infection by GFP expression, were systematically reduced in the absence of pre-DC by 22.5% with the R5 tropic virus (n=8, Figure 2a). The reduction was even more pronounced with the X4 tropic virus: an average decrease of 53.3% of DC infection in the absence of pre-DC (Figure 2b). These data indicated that despite pre-DC low representation among blood DC, this cell population has the potential to contribute to HIV-1 spreading due to its better capacity to capture HIV-1 and to get efficiently infected.

**Figure 2.**
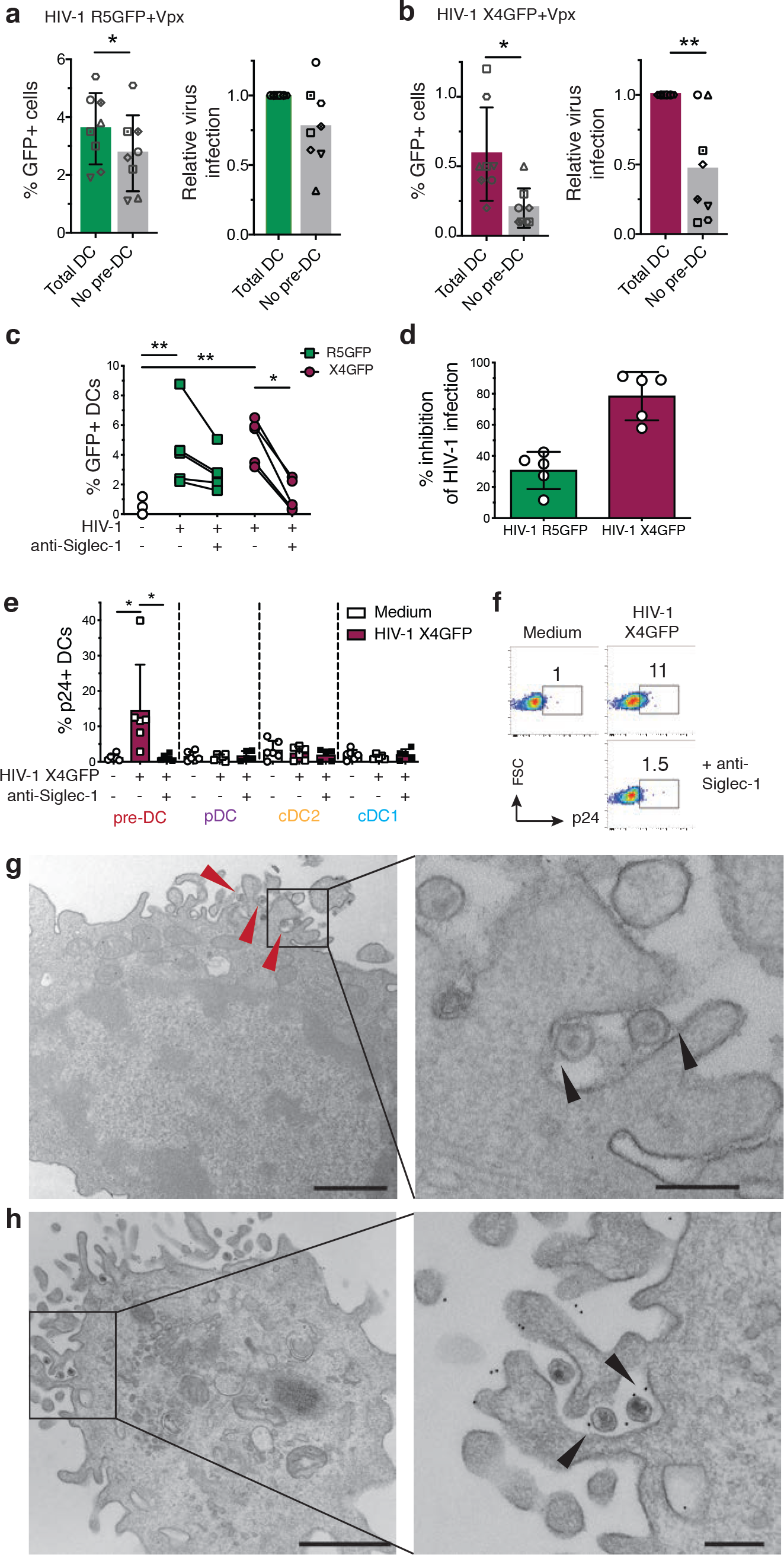
Pre-DC preferential infection is associated with Siglec-1 mediated capture of HIV-1. **a,** Quantification of absolute percentages (left panel) and relative (right panel) infection of total DC (CD45^+^Lin(CD3/CD14/CD16/CD19/CD20/CD34)^−^HLA-DR^+^) or total DC depleted of pre-DC (CD33^+^CD45RA^int^CD123^+^Axl^+^) exposed to HIV-1 R5GFP+Vpx or **b,** to HIV-1 X4GFP+Vpx. GFP expression was analysed 48h post-infection. **c,** GFP expression in sorted pre-DC pre-incubated with an anti-Siglec-1 mAb or not before being infected for 48 h with HIV-1 X4GFP or R5GFP, as in (Fig. 1c, d and e). Individual donors are displayed. **d,** Quantification of the inhibition of the HIV-1 infection by pre-incubation with an anti-Siglec-1 mAb. Data from the experiments depicted in (c) were processed. **e,** Quantification of HIV-1 capture by the four DC subsets. Sorted DCs were pre-incubated for 30 min with an anti-Siglec-1 mAb or not, then exposed or not to HIV-1 X4GFP for 2 h at 37°C, washed, stained for p24 to detect bound particles and assayed by flow cytometry (HIV-1 X4GFP viral particles are not GFP^+^ by themselves), n=5 or 6 independent donors combined in 5 experiments. Individual donors are displayed with bars representing mean ± S.D. **f,** Representative dot plot analysis of HIV-1 capture by pre-DC performed as in (e). **g,** Ultrastrucutral analysis of pre-DC exposed for 2 h to HIV-1 R5GFP, washed and processed for electron microscopy (EM). An epon section is presented, with a magnified view on the right panels. Arrowheads point to virions captured by pre-DC. On the magnified view, arrowheads point to protein linking the virions to the plasma membrane of the cells. Bar scale: 2µm (left image) and 0.5µm for the magnified view. **h,** Immuno-EM analysis of pre-DC exposed to HIV-1 R5GFP for 2 h fixed and stained for Siglec-1 with a mAb revealed by protein A gold coupled to 5 nm gold particles before embedding (see methods). An epon section is presented, with a magnified view. Arrowheads point to Siglec-1 specific staining at the interface between virions and plasma membrane invaginations. Bar scale: 2µm (left image) and 0.5µm for the magnified view.

To evaluate the potential role of Siglec-1 on pre-DC infection, pre-DC were exposed to Siglec-1 specific antibody prior to infection. This blockade prevented HIV-1 infection of pre-DC by R5-tropic viruses to some extent (35% inhibition) but more extensively for X4-tropic ones (roughly 85% inhibition, Figures 2c and 2d). Together these data suggested that Siglec-1 expression may endow pre-DC to better capture HIV-1 particles and stabilize their interaction leading to better rates of infection.

Thus, we first compared the various DC subsets for their capacity to capture HIV-1. Sorted DC populations were exposed to HIV-1 X4 for 2 h at 37°C, washed, and stained for p24 before analysis by flow cytometry. Among the four subsets, only pre-DC were able to capture HIV-1 (Figure 2e). The Siglec-1 involvement in the capture was evaluated by pre-exposure of pre-DC, as compared to the other DC subsets, to an anti-Siglec-1 antibody, which totally prevented capture of HIV-1 (Figures 2e and 2f). Moreover, EM ultrastructural analysis of pre-DC exposed to HIV-1 for 2 h revealed the presence of HIV-1 particles tightly bound to micro-villi at the pre-DC plasma membrane (Figure 2g). Analysis of ultrathin immunolabeled sections revealed that Siglec-1 protein was localized at the ultrastructural level in the immediate vicinity of both viral particles and the plasma membrane (Figure 2h). Given the very large size of the highly glycosylated Siglec-1 protein (1709 aa), these images are highly suggestive of Siglec-1 bridging the plasma membrane of pre-DC to HIV-1 particles. Thus, among blood DC, pre-DC are uniquely equipped with Siglec-1 that mediates efficient HIV-1 capture and infection.

### TLR activation of pre-DC induces RAB15 upregulation and prevents HIV-1 fusion and infection

To identify other factors that could explain the preferential susceptibility of pre-DC to HIV-1 as compared to other DC subsets, we first evaluated the expression levels of HIV-1 co-receptors on the various DC subsets. However, CCR5 and CXCR4 expression levels were similar among pre-DC, cDC2 and pDC (Figures 3a and 3b). The molecular basis for the resistance of cDC1 and pDC to HIV-1 infection at steady state has been recently attributed to their higher expression of the small GTPase RAB15^14^. Accordingly, we found that both HIV-1-suceptible subsets; pre-DC and cDC2 exhibited similar low levels of RAB15 mRNA, while these levels were higher in HIV-1-resistant cells i.e. pDC and cDC1 (Figure 3c). Since RAB15 expression has been proposed to inhibit HIV-1 infection by blocking viral fusion^14^, we measured viral fusion using a BlaM-Vpr assay in the 4 DC subsets purified *ex vivo* (Figures 3d and 3e). Strikingly, pre-DC and cDC2 exhibited a high capacity to fuse with HIV-1 (close to 80% of the cells) while the two HIV-1 resistant DC subsets; pDC and cDC1 had lower capacity (40% and less than 20%, respectively). Thus, rates of viral fusion were higher than rates of infection for each of the 4 subsets purified *ex vivo*.

**Figure 3.**
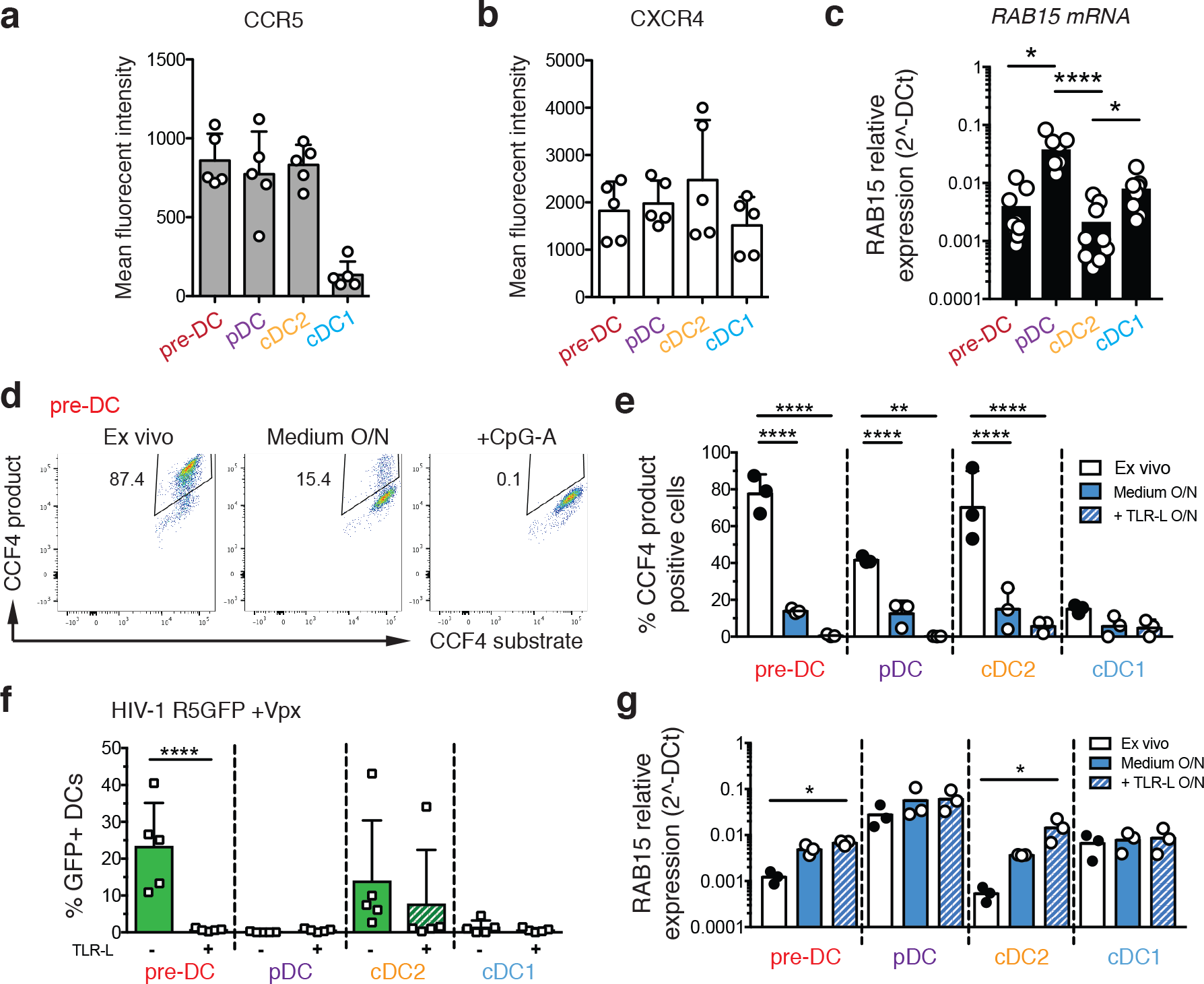
Pre-DC high capacity to fuse with HIV-1 and to get infected is lost upon activation and is associated with RAB15 expression levels. **a,** Surface expression levels of CCR5 and **b,** CXCR4 on the four freshly sorted DC subsets determined by flow cytometry, n=5 independent donors combined in 2 experiments. Individual donors are displayed with bars representing mean ± S.D. **c,** RAB15 mRNA expression levels among the four DC subsets freshly sorted, measured by qPCR, n=9 independent donors combined in 3 experiments. Individual donors are displayed with bars representing mean ± S.D. **d,** Representative dot plot of viral fusion revealed by CCF4 fluorescence in sorted pre-DC after infection with HIV-1 R5GFP containing a BlaM-Vpr fusion protein directly *ex vivo* or after an overnight culture with medium alone or in the presence of CpG-A. Fluorescence of the CCF4 product indicates viral fusion with target cells as a result of cleavage of the cell-loaded CCF4 substrate by the virus-contained BlaM. **e,** Quantification of viral fusion revealed by CCF4 fluorescence after infection with HIV-1 R5GFP containing a BlaM-Vpr fusion protein directly ex vivo or after an overnight culture with medium alone or in the presence of TLR-L (CpG-A for pre-DC and pDC, or CL264 for cDC2 and cDC1) for the four sorted DC populations, n=3 donors combined from two independent experiments. **f,** Quantification of DC infection with HIV R5GFP+Vpx directly *ex vivo* or following overnight (O/N) incubation with TLR-ligands (CpG-A for pre-DC and pDC; CL064 for cDC2 and cDC1). GFP expression was measured at 48h post-infection, n=5 from 3 independent experiments. Individual donors are displayed with bars representing mean ± S.D. **g,** RAB15 mRNA levels among the four DC subsets freshly sorted, measured by pPCR in DC subsets, *ex vivo* or after an overnight culture in medium alone or supplemented with TLR-L as in (E), n=3.

Given that systemic immune activation occurs in both acute and chronic HIV infection in part due to compromised integrity of gut epithelium^28^, we asked whether TLR-mediated activation on DC could impact their susceptibility to viral fusion. Strikingly, while overnight culture already substantially reduced the fusion rates, overnight TLR activation induced a total block in HIV-1 fusion for all DC subsets (Figure 3e). Indeed, overnight TLR activation abolished infection of pre-DC and of cDC2 (in 4 out of 5 patients), while pDC and cDC1 remained resistant to HIV-1 (Figure 3f). Accordingly, overnight culture induced the expression of RAB15 mRNA in both pre-DC and cDC2, which was further increased following TLR activation.

We concluded that TLR activation completely switches pre-DC phenotype from highly susceptible to total resistance to HIV-1 infection. This phenotypic change is accompanied by an increased expression of RAB15 and a complete resistance to HIV-1 fusion.

### HIV-1 replicates in pre-DC apparently internal compartments

Since HIV-1 can efficiently fuse with both pre-DC and cDC2, we next examined the infected cells at the intracellular level to follow the late steps of the viral cycle. Pre-DC and cDC2 infected for 48 h with HIV-1 R5 + Vpx were analyzed by electron microscopy (EM) (Figures 4a, 4b and S3). Strikingly, pre-DC exhibited typical Virus-Containing Compartments (VCC) similar to the ones observed in HIV-1-infected MDM^29–31^. Viral budding profiles were mostly absent from the plasma membrane but were clearly observed at the limiting membrane of the compartment, indicating *de novo* production of viral particles. These profiles exhibited a dark molecular coat typical of Gag precursor assembly^32,33^, that budded towards the lumen of the compartments in which numerous immature and mature viral particles accumulated (Figure S3a). A specific feature of the VCC from macrophages is their accessibility to the external medium that can be evaluated by adding ruthenium red, a membrane impermeant dye, during fixation^34,35^. Ruthenium red was visible in pre-DC VCC as a dark and fuzzy shade at the limiting membrane of the VCC and on the viral membranes (Figure 4a and S3a), demonstrating that these compartments are connected to the external medium. Interestingly, similar internal compartments preexist to the infection in MDM^30,36^ and are hijacked by HIV-1 for its assembly, leading to their conversion into VCC.

**Figure 4.**
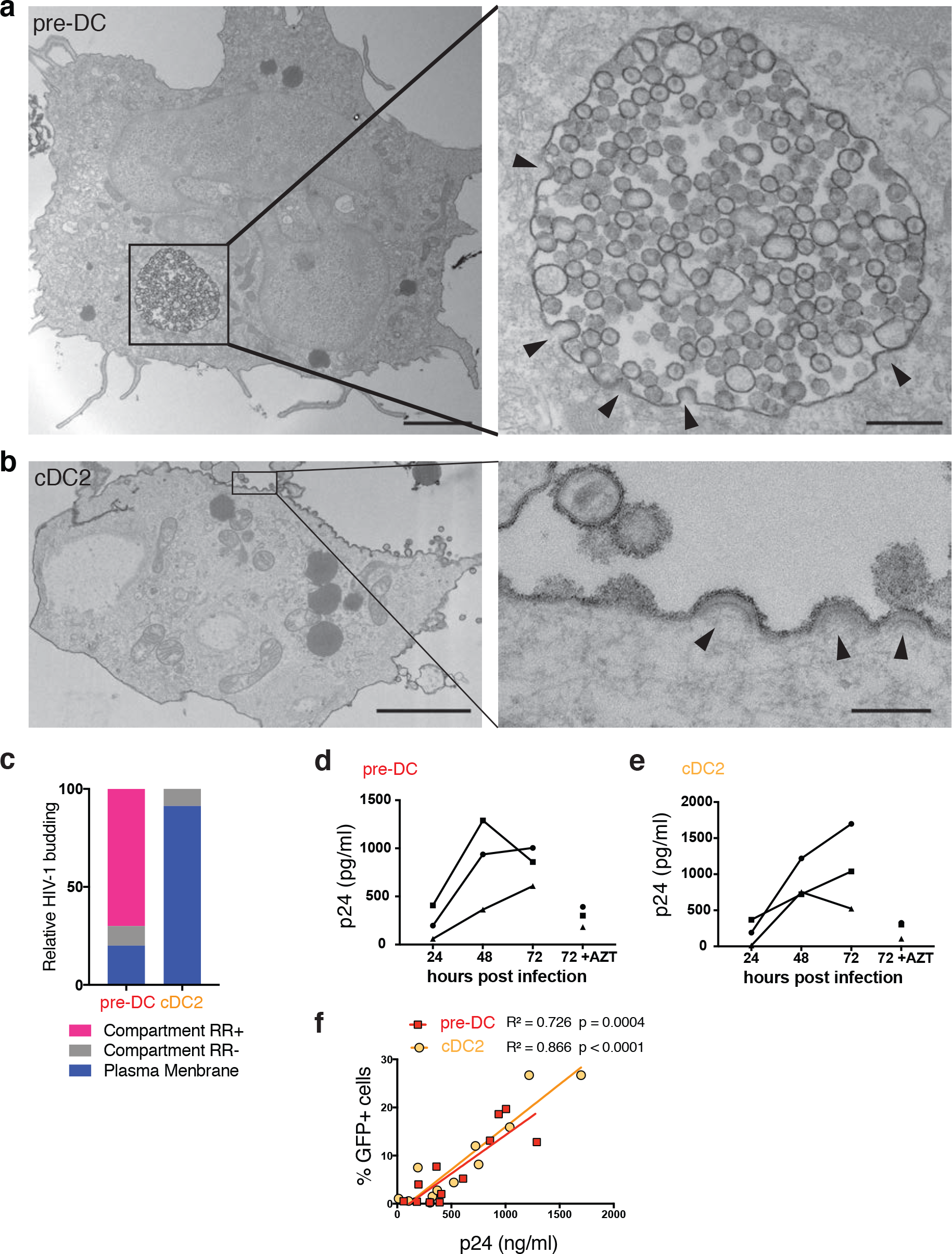
HIV-1-infected pre-DC, but not cDC2, produce new virions in apparently intracellular compartments. **a,** Epon sections from sorted pre-DC and **b,** cDC2 infected with HIV-1 R5GFP+Vpx for 48 h and analysed by EM. Cells were fixed in the presence of ruthenium red to label surface and surface-connected compartments. Arrowheads point to nascent viral buds. Bar scale, 2µm (left images) and 0,15µl for magnified views. **c,** Quantification of the location of viral budding profiles observed in HIV-1 infected pre-DC (n=11) and cDC2 (n=22). RR, ruthenium red. Of note, some internal compartments containing viruses were observed in infected cDC2, they were however unlabeled by RR and did not exhibit viral budding profiles at their limiting membranes. Rather than VCC, they probably represent endosomes having internalized viral particles secreted by neighboring cells. **d,** Kinetics of production of p24 by sorted pre-DC and **e,** cDC2 infected with HIV-1 R5GFP+Vpx measured by CBA (see methods). The RT inhibitor AZT was used as a control for the viral input remaining at the end of the assay, n=3 independent donors combined in 2 experiments. Individual donors are displayed. **f,** Correlation between the levels of infection of pre-DC (red) and cDC2 (yellow) measured by GFP expression and the quantification of p24 in supernatants as in (d and e).

In sharp contrast, infected cDC2 were free of VCC and viral budding only occurred in a polarized manner at the plasma membrane (Figures 4b and S3b), as previously observed in infected T lymphocytes and cell lines^37^. In cDC2, ruthenium red only stained the plasma membrane and released viral particles. Quantification of compartments accessible to ruthenium red that exhibited viral buds and particles confirmed the striking differences between the two subsets (Figure 3c). Thus, pre-DC and cDC2, although related through their precursor-to-progeny relationship as pre-DC can give rise to cDC2^6,7^, exhibit different susceptibility to infection and completely different locations of viral assembly.

The EM profiles indicated that active viral replication can take place in both cell types. We compared their production after pre-DC and cDC2 exposure to HIV-1 R5GFP complemented with Vpx. Both infected DC subsets released in their culture supernatant increasing amounts of p24 overtime (Figures 4d and 4e) that correlated with the rates of infection (Figure 4f), indicating that HIV-1 can replicate in both subsets.

### Pre-DC transmit HIV-1 to CD4^+^T cells in *cis* or *trans* depending on their activation state

The infectious capacity of the viral particles produced by HIV-1-infected pre-DC was evaluated on primary activated CD4^+^ T lymphocytes (Figure 5a). In line with their preferential infection with X4-tropic virus, HIV-1-infected pre-DC were the most efficient subset in transmitting the infection in *cis* to activated primary CD4^+^ T cells (Figures 5b and 5c). These results establish that the viral progeny of infected pre-DC is infectious and thus may contribute to viral spreading.

**Figure 5.**
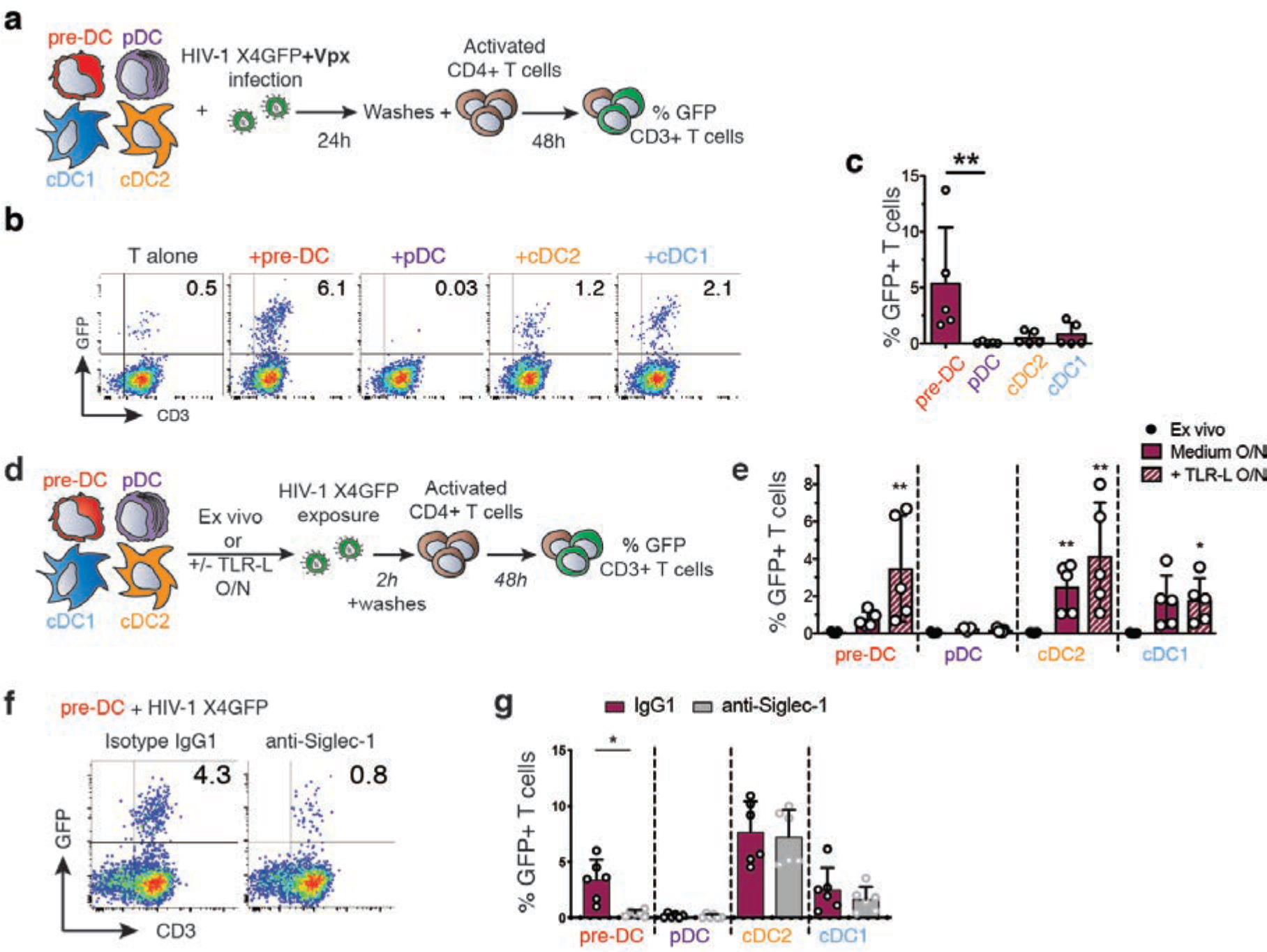
Infected pre-DC and activated non-infected pre-DC can both transmit HIV-1 to activated CD4+ T cells. **a,** *Cis*-infection of T cells, outline of the experiment. **b,** GFP expression in CD3^+^T cells co-cultured with HIV-1-infected DC subsets or not, as explained in (a). Representative dot plots are presented. **c,** Quantification of GFP expression in CD3^+^T cells as in (b), n=5 independent donors combined from 2 experiments. Individual donors are displayed with bars representing mean ± S.D. **d,** *Trans*-infection of T cells, outline of the experiment. **e,** GFP expression in activated T cells co-cultured directly with *ex vivo* DC or following overnight culture in medium or in the presence of TLR-L (CpG-A for pre-DC and pDC, CL264 for cDC). Prior to CD4+T cell co-culture, DC populations were exposed to HIV-1 X4GFP (without Vpx) for 2 h and washed. Quantification of GFP^+^ T cells (CD3^+^) was performed by flow cytometry after a 48h co-culture, n=5 independent donors combined in two experiments. Individual donors are displayed with bars representing mean ± S.D. **f,** GFP expression by CD3^+^T cells. TLR-activated pre-DC were pre-incubated with anti-Siglec-1 antibody or with an isotype control antibody (IgG1) before performing the *trans-*infection assay as in (e). **g,** Quantification as in (f) with the indicated DC populations. (n=6 independent donors combined in 3 experiments). Individual donors are displayed with bars representing mean ± S.D.

Given that TLR stimulation induced a state of resistance to HIV-1 infection, we evaluated the capacity of DC populations to perform HIV-1 *trans-*infection of T cells. During *trans*-infection DC capture HIV-1 via Siglec-1 but importantly, do not get infected and efficiently transfer the virus to T cells as shown *in vitro* with LPS-activated MDDC^16,18,20^. Hence, we examined whether pre-DC activated or not were able to support HIV-1 infection of T cells in *trans*, compared to the other DC subsets (Figure 5d). First, none of the *ex-vivo* sorted DC populations, including pre-DC, was able to transmit HIV-1 to activated CD4^+^ T cells (Figure 5e). However, after overnight culture, pre-DC, cDC1 and cDC2 performed *trans-*infection to CD4^+^ T cells, which was further increased upon TLR-activation (Figure 5e). Similarly, MDDC cultured overnight in the presence or absence of LPS performed *trans*-infection of T cells (Figure S4a). Thus, following TLR activation, both pre-DC and cDC2 performed HIV-1 *trans-*infection to CD4^+^ T cells to similar levels, while cDC1 did poorly and pDC did not (Figure 5e). Finally, pre-incubation of TLR-activated DC subsets with anti-Siglec-1 antibody strongly inhibited pre-DC but not cDC2 capacity to perform *trans-*infection (Figures 5f and 5g), and reduced the one of MDDCs exposed to LPS (Figure S4b). We concluded that TLR-activated pre-DC can perform genuine *trans-*infection, capturing HIV-1 without being infected (Figure 3f) and transmitting the virus efficiently to activated CD4^+^ T cells. The fact that *ex-vivo* sorted unstimulated pre-DC are unable to perform *trans-*infection to T cells despite their high levels of surface Siglec-1, indicates that activation of pre-DC induces the acquisition of this capacity through a still unknown mechanism.

## Discussion

The recent identification of the human blood pre-DC population was based on single cell data analysis using unsupervised clustering approaches^6,7^. Here we show that pre-DC exhibit a unique set of properties regarding HIV-1 that are linked to their constitutive Siglec-1 expression (Figure S5). Indeed we show first that Siglec-1 is pivotal for the unique capacity of pre-DC to capture HIV-1. In addition, among freshly sorted blood DC subsets, pre-DC stands out as the most susceptible to infection by both types of HIV-1; CCR5-tropic viruses but also surprisingly, CXCR4-tropic ones. This type of virus is unable to infect any other DC subtype even when the SAMHD1 restriction factor is counteracted (^14^ and Figure 1). The high capacity of binding HIV-1 via Siglec-1 indicate that these cells could act as flypaper for the virus in the blood stream and maybe in tissues as well. Despite their low frequency among blood DC, pre-DC were able to compete out other DC and get preferentially infected, especially with CXCR4 viruses.

Stabilization of the interaction between HIV-1 and pre-DC via Siglec-1 may increase the probability of the Env-mediated fusion to occur, a rather slow process^38,39^. It may also account for pre-DC preferential susceptibility to X4 viruses as compared to other blood DC populations that do not express Siglec-1, despite their common low levels of expression of the co-receptor CXCR4. Regarding CCR5-tropic viruses, we observed that both pre-DC and cDC2 had a high capacity to fuse with this type of virus (around 80%). However, cDC2 exhibited a much lower infection rate than pre-DC, even when the SAMHD1 restriction factor was counteracted (roughly 20% versus 50%). On the same line, while pDC are totally resistant to infection by R5 HIV-1, 40% of them were susceptible to viral fusion as judged using the BLaM-Vpr assay. The discrepancies between the rates of fusion and infection, even when SAMHD1 was counteracted, suggest that additional unknown mechanisms are at work in cDC2 and pDC.

RAB15 high expression has been associated to resistance to viral fusion of cDC1 and thus to infection^14^. We confirmed and extended these data documenting high levels of RAB15 transcripts in pDC and cDC1 and low levels in cDC2 and pre-DC at the steady state. Importantly, we further established that TLR activation upregulates these levels in all DC subsets and confers to all of them full resistance to fusion and infection. More work is needed to understand the molecular mechanism involved but the comparison of the various DC subsets may represent a valuable approach for this purpose.

Our data also indicate that surface Siglec-1 on pre-DC facilitates pre-DC infection and further, transmission to activated T cells in *cis*. Infection leads to viral production and accumulation in apparently intracellular compartments termed VCC. Of note, Siglec-1 is also present at the VCC limiting membrane of infected MDM^40^ and thus in the HIV-1 particles they produce^41^. Therefore, Siglec-1 expression by pre-DC might be implicated in HIV-1 assembly at the VCC level. VCC have been initially observed in tissues macrophages from infected patients in a variety of tissues, see ^32^, and were then essentially studied in *in vitro* derived and infected MDM^31^. Importantly, a recent study identified urethral macrophages from individual under cART containing infectious particles in VCC, suggesting that such tissue macrophages can represent a viral reservoir^42^. In contrast, HIV assembly at the plasma membrane, as observed here in primary cDC2 for the first time, has been largely documented in activated CD4+ T cells. The different location of viral assembly between pre-DC and cDC2 is compelling since pre-DC can give rise to cDC2. Comparison between the two DC subtypes should help to approach the molecular basis for VCC formation and the potential role of Siglec-1 in this process, as previously proposed^43^.

When activated, pre-DC are no longer susceptible to infection but can perform efficient *trans-*infection to CD4^+^ T cells, again in a fully Siglec-1-dependent manner. We thus confirm with primary pre-DC what has been established with LPS-stimulated MDDC, i.e. the existence of this mode of transmission to T cells via non-infected DC. The fact that *ex-vivo* purified pre-DC were unable to perform HIV-1 trans-infection suggests that DC activation is necessary. In contrast to a previous study performed with LPS-stimulated cDC2^17^, we observed that TLR7-activated cDC2 performed trans-infection independently of Siglec-1. The impact of different TLR stimulations of DC on the HIV-1 *trans*-infection process remains to be established.

How pre-DC integrate in the highly organized share of work between the various DC subsets to face or contribute to the HIV infection remains to be established. Recent work of Silvin et al. points out that cDC2 get infected by HIV-1, whereas cDC1 resist entering in a resistant state that allows them to cross-present viral antigens from dead infected cDC2 cells^14^. As pre-DC can secrete IL-12 and support mixed-lymphocyte reactions^6,7^, it will be of interest to determine whether and how pre-DC possess the hybrid capacity of both cDC1 and cDC2 to initiate an anti-HIV-1 response.

Interestingly, Siglec-1^+^ macrophages from the spleen marginal zone have been shown in the VSV mouse model to support a locally restricted enforced viral replication by down-modulating type I IFN responsiveness thus providing a source of viral antigen required to elicit a protective immune response^44^. Moreover, Siglec-1+ macrophages have been recently shown to be able to transfer viral antigens to cDC1 which in turn cross-prime CD8+ T cell responses. Importantly, this antigen transfer was mediated by Siglec-1 on the macrophages interacting with sialylated proteins present on cDC1 cell surface. Thus, Siglec-1 expression may also confer pre-DC the capacity to recruit and instruct the help of cross-presenting cDC1. Future studies will certainly address this important hypothesis, since manipulation of pre-DC response or protection against HIV-1 may represent new approaches to promote a better immune control of the infection.

## Methods

### Isolation of cells

Peripheral blood mononuclear cells were isolated from buffy coats from healthy human donors (approved by the Institut National de la Santé et de la Recherche Médicale ethics committee and the Health Sciences Authorities (Singapore)) with Ficoll-Paque PLUS (GE). Total blood DCs were enriched with EasySep human pan-DC pre-enrichment kit (Stemcell Technologies). DC enriched fraction were stained with antibodies specific for HLA-DR APCeFluor780, CD1c PerCPeFluor710 (eBioscience), CD123 Viogreen, CD141 Vioblue (Miltenyi), CD33 APC, CD45RA PE (BD) and with a cocktail of antibodies against lineage markers CD19 (Miltenyi), CD3, CD14, CD16 and CD34 (BD) in the FITC channel. Alternatively, DC enriched fraction were stained with HLA-DR APCeFluor780, CD1c PerCPeFluor710, CD123 Viogreen, CD45RA Vioblue (Miltenyi), CD33 PE-CF594, Clec-9A PE (BD) with lineage markers in the FITC channel. Pre-DC were sorted as Lin^-^ HLADR^+^ CD33^int^ CD45RA^int^ CD123^+^. pDCs were sorted as Lin^-^ HLADR^+^ CD33^-^ CD45RA^+^ CD123^+^. cDC2 were sorted as Lin-HLADR^+^ CD33^+^ CD45RA^-^ CD1c^+^. cDC1 were sorted as Lin^-^ HLADR^+^ CD33^+^ CD45RA^-^ CD141^+^/Clec9A^+^.

For pre-DC depletion, DC enriched fraction were stained with antibodies specific for HLA-DR APCeFluor780 with a cocktail of antibodies against lineage markers CD19, CD3, CD14, CD16 and CD34 in the FITC channel or with the addition of CD123 Viogreen, Siglec-1 PE-Vio770 (Miltenyi), Axl PE (Clone #108724, R&D Systems). Total DC were sorted as Lin^-^ HLADR^+^. Alternatively, Pre-DC (Lin^-^ HLADR^+^ CD123^+^ Siglec-1^+^ Axl^+^) were excluded from Lin^-^ HLADR^+^.

All cells were sorted on a FACSAria (BD) using Diva software (BD) in 5mL polypropylene round-bottom tubes containing 1mL X-VIVO-15 media (Lonza BE04-418F). Post-sort cell purity after gating on live cells by FCS/SSC was routinely between 92 and 99%.

CD4^+^ T cells were isolated from PBMC by negative selection using CD4^+^ T Cell Isolation Kit (Miltenyi 130-096-533). CD14^+^ monocytes were obtained by positive selection from PBMC using CD14 Microbeads (Miltenyi 130-050-20).

### DC phenotype by flow cytometry

Enriched DC obtained as described were stained with HLA-DR APCeFluor780, CD1c PerCPeFluor710, CD123 Viogreen, CD45RA Vioblue, CD33 PE-CF594, CD141 BV711 (BD), Axl PE, Siglec-1 PE-Vio770 with lineage markers in the FITC channel. Cells were acquired on a FACS Verse and data analyzed using FlowJo 10 (Tree Star Inc.). After compensation, tSNE representation of gated Lin^-^ HLADR^+^ cells were generated. Siglec-1 and Axl expression were evaluated on gated pre-DC, pDC, cDC1 and cDC2 as described.

Alternatively, PBMCs were stained as described previously^6^ with DC lineage markers in addition to Siglec-1 (BD), CXCR4 (Biolegend) and CCCR5 (BD) and analysed by flow cytometry. FCS files compensated for spillover between channels were exported using FlowJo v10 (Tree Star Inc.) and CXCR4 and CCR5 expression were measured on gated pre-DC, pDC, cDC1 and cDC2 as described.

### Cell Culture

For lentiviral infections of sorted DC, cells were cultured in X-VIVO-15 complemented with Penicillin/Streptomycin (Thermo Fisher 10378-016). For capture experiments, DCs were cultured in RPMI 1640 medium, GlutaMAX (Thermo Fisher 61870-010) complemented with FBS 10% (Thermo Fisher 10270-106) and Penicillin/Streptomycin.

T cells were cultured in RPMI 1640 medium, GlutaMAX complemented with FBS 10% and Penicillin/Streptomycin at 10^6^ cells/mL in the presence of 5µg/mL PHA (Lectin from Phaseolus vulgaris Leucoagglutinin; Sigma L2769) and 50U/mL of IL-2 (eBioscience). On day 2 of culture, cells were washed and additionally cultured with IL-2.

Purified CD14^+^ cells were cultured at 0.8 10^6^ cells/mL in complete RPMI supplemented with GM-CSF at 100ng/mL and IL-4 (Miltenyi) at 50ng/mL for 4 days to obtain monocyte-derived DC (MDDC).

HEK293FT cells were cultured in DMEM medium, GlutaMAX (Thermo Fisher 61965-026) complemented with FBS 10% and Penicillin/Streptomycin. GHOSTX4R5 cells were cultured in DMEM medium, GlutaMAX complemented with FBS 10% and Penicillin/Streptomycin.

All cells were cultured at 37°C with 5% CO_2_ atmosphere.

### Plasmids

The proviral plasmid HIV-1 R5GFP was derived from of NL4-3 with BaL env, Δnef and GFP in nef^45,46^. HIV-1 X4GFP corresponded to NL4-3, Δnef, GFP in nef. pIRES2EGFP-VPXanyVPR plasmid was used to complement some of the viral preparations with Vpx^47^ and pMM310 (BlaM-Vpr) to complement the viral preparations with BlaM to measure the fusion of virions^48^.

### HIV-1 production and titration

Viral particles were produced by transfection of HEK293FT cells in 6-well plates with 3µg DNA and 8µl TransIT-293 (Mirus Bio) per well. For HIV-1 R5GFP and HIV-1 X4GFP, 3µg of HIV plasmid were used. For HIV-1 virus containing Vpx and BlaM, 0.75µg pIRES2EGFP-VPXanyVPR or pMM310 respectively and 2.25µg HIV plasmid were used. 16h after transfection, media was removed, and fresh X-VIVO or RPMI medium was added. Viral supernatants were harvested 36h later, filtered at 0.45µM, used freshly or aliquoted and frozen at −80°C. Viral titers were determined on GHOST X4R5 cells as described^49^ and multiplicity of infection (MOI) ranged from 0.04 to 1.99, mean=0.99, S.D.=0.49 for DC infection displayed in Figure 1.

### HIV infection of DCs and stimulations

Sorted cells were pelleted and resuspended in complete X-VIVO or RPMI media at 0.4 10^6^ cells/mL and 50µl were seeded in round bottom 96-well plates. In some experiments anti-Siglec-1 mAb (clone 7-239) or mIgG1 isotype control (Miltenyi) were added at 20µg/mL and cells incubated for 30 min. at 37°C before adding the virus. Azidothymidine (AZT; Sigma) was used at 25 µM final concentration, and nevirapin (Sigma) was used at 5 µM. CpG-A (ODN2216, Invivogen) was used at 5µg/mL, CL264 (Invivogen) at 10µg/mL. For infections, 150µl of media or dilutions (150µl or 50µl) of viral supernatants were added. In some cases, when mentioned, infections were spinoculated for 2h at 800g 25°C.

For total DC or pre-DC depleted DC, cells were were resuspended at 2.10^6^ cell/ml and 2.10^5^icells were cultured in the presence of 450µl mock medium or HIV-1 R5GFP viruses in X-vivo.

48hr after infection, cell culture supernatants were harvested and cells were fixed in 4% paraformaldehyde (PFA; Electron Microscopy Sciences) in PBS prior to analysis on a FACSVerse flow cytometer (BD). Data were analyzed using FlowJo v10 and Prism v7 for Mac (GraphPad).

### Electron microscopy

For routine embedding into Epon, cells were fixed in 2.5% glutaraldehyde in 0.1M Na-Cacodylate buffer pH 7.4 for 1h, postfixed for 1h with 1% buffered osmium tetroxide, dehydrated in a graded series of ethanol solution, and then embedded in epoxy resin as described^35^. Electron micrographs were acquired on a Tecnai Spirit electron microscope (FEI, Eindhoven, The Netherlands) equipped with a 4k CCD camera (EMSIS GmbH, Münster, Germany). For pre-embedding immunogold labeling, cells were fixed in 4% paraformaldehyde for 1h at room temperature, quenched in PBS-Glycine and blocked with 1% BSA-PBS. Primary antibody anti-Siglec-1 mAb (Miltenyi) was diluted to 1:50 in 1% BSA-PBS and incubated with cells for 1h. Cells were extensively washed prior to addition of protein A gold (EM lab, Utrecht University), and routine embedding in Epon.

For Rethunium Red (RR) stain, cells were fixed with 2.5% glutaraldehyde in 0.1M ice cold Na-Cacodylate buffer (pH 7.4) containing 0.15% RR overnight at 4°C. Cells were then washed with 0.1M Na-Cacodylate buffer (pH 7.2), post-fixed with 1.3% OsO4 in the same buffer containing 0.15% RR for 2 h at room temperature dehydrated in a graded series of ethanol solution, and then embedded in epoxy resin.

### HIV capture and trans-infection of activated CD4^+^ T cells

Sorted cells were pelleted and resuspended in complete X-vivo media at 0.4 10^6^ cells/mL and 50µl were seeded in round bottom 96-well plates. In some experiments anti-Siglec-1 mAb or mIgG1 isotype control were added at 20µg/mL and cells incubated for 30 min. at 37°C before adding the virus. HIV-1 X4GFP was added onto the cells (150µl/well of HEK293FT culture supernatant) and incubated for 2h at 37°C. Cells were washed extensively and fixed in 4% PFA in PBS. p24 staining was performed using KC-57 RD1 mAb (Bekman Coulter, 6604667). For trans-infection experiment, sorted DC were washed extensively after the 2h-culture with HIV-1 X4GFP and activated CD4^+^ T cells were added at a ratio 1:1. Alternatively, CpG-A was added at 5µg/mL, CL264 was added at 10µg/mL onto DCs and cells incubated overnight before the addition of HIV-1 X4GFP. Cells were then washed and activated CD4+ T cells added. After 48h, cells were fixed in 2% PFA in PBS. Cells were then stained with PE-Cy7 anti-CD3 (BD) and analyzed on a FACS Verse (BD). Data were analyzed using FlowJo v10 and Prism v7 for Mac (GraphPad).

### HIV-1 fusion assay

DC infection was performed as described with 150µl BlaM-containing viral supernantant and cells were spinoculated for 2h at 800g, 25°C. Cells were additionally incubated 3hr at 37°C, washed 3 times with CO2 independent medium (Thermo Fisher 18045-054), loaded with CCF4-AM dye (dilution 1/400) in CO2 independent medium complemented with solution B (dilution 1/125) for 1hr at room temperature (RT). Cells were washed two times with CO2 independent medium and resuspended in 100µL of CO2 independent medium complemented with 10% FBS and 2.5 mM probenicid. Cells were incubated 12hr at RT, washed with CO2 independent medium and fixed with 1% PFA in PBS before acquisition on a FACS Verse. Data were analyzed using FlowJo v10 and Prism v7 for Mac.

### Microarray analysis^6^

Total RNA was isolated from FACS-sorted blood pre-DC and DC subsets using a RNeasy^®^ Micro kit (Qiagen). Total RNA integrity was assessed using an Agilent Bioanalyzer (Agilent) and the RNA Integrity Number (RIN) was calculated. All RNA samples had a RIN ≥7.1. Biotinylated cRNA was prepared using an Epicentre TargetAmp™ 2-Round Biotin-aRNA Amplification Kit 3.0 according to the manufacturer's instructions, using 500 pg of total RNA starting material. Hybridization of the cRNA was performed on an Illumina Human-HT12 Version 4 chip set (Illumina). Microrarray data were exported from GenomeStudio (Illumina) without background subtraction. Probes with detection P-values > 0.05 were considered as not being detected in the sample, and were filtered out. Expression values for the remaining probes were log2 transformed and quantile normalized. Microarray data set is deposited in the Gene Expression Omnibus under accession number GSE80171.

### RAB15 gene expression

4.10^5^ sorted DC were used *ex-vivo* or treated as described. Cells were lysed, RNA was extracted using RNeasy micro kit (Qiagen) and reverse transcription was performed using a high-capacity cDNA reverse transcription kit (Applied Biosystems) by following the manufacturer's instructions. Real-time quantitative PCR was performed using SYBR Green I Master (Roche) with the following primers (Eurogentec): RAB15-forward: 5’-CAGCAGCTGGCGAAGGAG-3’ and RAB15-reverse: 5’-GTTGGTGCAGGCACTTCTTTC-3’. RPS18 gene was used as a house-keeping gene with the following primers: RPS18-forward: 5’-CTGCCATTAAGGGTGTGG-3’, RPS18-reverse: 5’-TCAATGTCTGCTTTCCTCAAC-3’. The relative quantity of RAB15 mRNAs was calculated between RAB15 RNA Cp and RPS18 Cp by the 2-∆Ct method.

### Measure of p24 by Cytometric Bead Assay (CBA)

In house p24 CBA assay was used to quantify p24 concentration in the supernatant of HIV-1 infected DC. Dilutions of virus-like particles with known concentration of p24 were used to establish a standard curve. 300 beads/sample were acquired on a FACS Verse. Data were analyzed using FlowJo v10 and Prism v7 for Mac.

### Statistical analysis

Data were analyzed using Prism v7 for Mac (GraphPad). Friedman test was used followed by Dunn’s multiple comparisons tests and a p value lower than 0.05 was considered as significant (*p<0.05; **p<0.01; ***p<0.001 and ****p<0.0001).

## Author contributions

NR performed most of the experiments with the help of EGM, FB and CAD. MJ performed all the EM analyses. AS provided initial advice and help for sorting strategies. FG provided initial suggestion to study pre-DC and HIV-1 relationship and advice all along the study. NR and PB designed the experiments and analyzed the data. NR built the figures. PB wrote the manuscript with the contribution of NR, CAD and FG.

## Acknowledgments

We acknowledge Nicolas Manel at Institut Curie for fruitful discussions, reagents and critical reading of the manuscript. We also thank Ana-Maria Lennon-Dumenil, François-Xavier Gobert, Julie Helft, Constance Delaugerre and Sebastian Amigorena for discussions. We acknowledge the PICT-IBiSA for access to the imaging facilities and the flow cytometry facilities at Institut Curie and Institut Cochin (CYBIO). Microarray data set is deposited in the Gene Expression Omnibus under accession number GSE80171.

## References

1. Hladik, F. & McElrath, M. J. Setting the stage: host invasion by HIV. 8, 447–457 (2008).

2. Wu, L. & KewalRamani, V. N. Dendritic-cell interactions with HIV: infection and viral dissemination. 6, 859–868 (2006).

3. Cameron, P., Pope, M., Granelli-Piperno, A. & Steinman, R. M. Dendritic cells and the replication of HIV-1. J Leukoc Biol 59, 158–171 (1996).

4. Eickhoff, S. et al. Robust Anti-viral Immunity Requires Multiple Distinct T Cell-Dendritic Cell Interactions. Cell 162, 1322–1337 (2015).

5. Hor, J. L. et al. Spatiotemporally Distinct Interactions with Dendritic Cell Subsets Facilitates CD4+ and CD8+ T Cell Activation to Localized Viral Infection. Immunity 43, 554–565 (2015).

6. See, P. et al. Mapping the human DC lineage through the integration of high-dimensional techniques. Science 356, eaag3009 (2017).

7. Villani, A.-C. et al. Single-cell RNA-seq reveals new types of human blood dendritic cells, monocytes, and progenitors. Science 356, (2017).

8. Schlitzer, A., McGovern, N. & Ginhoux, F. Dendritic cells and monocyte-derived cells: Two complementary and integrated functional systems. Seminars in cell & developmental biology 41, 9–22 (2015).

9. Guilliams, M. et al. Unsupervised High-Dimensional Analysis Aligns Dendritic Cells across Tissues and Species. Immunity 45, 669–684 (2016).

10. Schlitzer, A. et al. Identification of cDC1- and cDC2-committed DC progenitors reveals early lineage priming at the common DC progenitor stage in the bone marrow. Nat Immunol 16, 718–728 (2015).

11. Breton, G. et al. Human dendritic cells (DCs) are derived from distinct circulating precursors that are precommitted to become CD1c +or CD141 +DCs. J Exp Med jem. 20161135–10 (2016). doi:10.1084/jem.20161135

12. Granelli-Piperno, A., Delgado, E., Finkel, V., Paxton, W. & Steinman, R. Immature dendritic cells selectively replicate macrophagetropic (M-tropic) human immunodeficiency virus type 1, while mature cells efficiently transmit both M- and T-tropic virus to T cells. J Virol 72, 2733–2737 (1998).

13. McIlroy, D. et al. Infection frequency of dendritic cells and CD4+ T lymphocytes in spleens of human immunodeficiency virus-positive patients. J Virol 69, 4737–4745 (1995).

14. Silvin, A. et al. Constitutive resistance to viral infection in human CD141+dendritic cells. Science Immunology 2, eaai8071 (2017).

15. Müller-Trutwin, M. & Hosmalin, A. Role for plasmacytoid dendritic cells in anti-HIV innate immunity. Immunol. Cell Biol. 83, 578–583 (2005).

16. Garcia, E. et al. HIV-1 trafficking to the dendritic cell-T-cell infectious synapse uses a pathway of tetraspanin sorting to the immunological synapse. Traffic 6, 488–501 (2005).

17. Izquierdo-Useros, N. et al. Siglec-1 is a novel dendritic cell receptor that mediates HIV-1 trans-infection through recognition of viral membrane gangliosides. PLoS Biol 10, e1001448 (2012).

18. Izquierdo-Useros, N. et al. Capture and transfer of HIV-1 particles by mature dendritic cells converges with the exosome-dissemination pathway. Blood 113, 2732–2741 (2009).

19. Puryear, W. B., Yu, X., Ramirez, N. P., Reinhard, B. M. & Gummuluru, S. HIV-1 incorporation of host-cell-derived glycosphingolipid GM3 allows for capture by mature dendritic cells. Proc Natl Acad Sci USA 109, 7475–7480 (2012).

20. Puryear, W. B. et al. Interferon-inducible mechanism of dendritic cell-mediated HIV-1 dissemination is dependent on Siglec-1/CD169. PLoS Pathog 9, e1003291 (2013).

21. Hrecka, K. et al. Vpx relieves inhibition of HIV-1 infection of macrophages mediated by the SAMHD1 protein. Nature 474, 658–661 (2011).

22. SAMHD1 is the dendritic- and myeloid-cell-specific HIV-1 restriction factor counteracted by Vpx. 474, 654–657 (2011).

23. Cribier, A., Descours, B., Valadão, A. L. C., Laguette, N. & Benkirane, M. Phosphorylation of SAMHD1 by cyclin A2/CDK1 regulates its restriction activity toward HIV-1. CellReports 3, 1036–1043 (2013).

24. Arnold, L. H. et al. Phospho-dependent Regulation of SAMHD1 Oligomerisation Couples Catalysis and Restriction. PLoS Pathog 11, e1005194 (2015).

25. SAMHD1 restricts the replication of human immunodeficiency virus type 1 by depleting the intracellular pool of deoxynucleoside triphosphates. 13, 223–228 (2012).

26. Bakri, Y. et al. The maturation of dendritic cells results in postintegration inhibition of HIV-1 replication. J Immunol 166, 3780–3788 (2001).

27. Cameron, P. U. et al. Dendritic cells exposed to human immunodeficiency virus type-1 transmit a vigorous cytopathic infection to CD4+ T cells. Science 257, 383–387 (1992).

28. Brenchley, J. M. et al. Microbial translocation is a cause of systemic immune activation in chronic HIV infection. 12, 1365–1371 (2006).

29. Tan, J. & Sattentau, Q. J. The HIV-1-containing macrophage compartment: a perfect cellular niche? Trends Microbiol 21, 405–412 (2013).

30. Mlcochova, P., Pelchen-Matthews, A. & Marsh, M. Organization and regulation of intracellular plasma membrane-connected HIV-1 assembly compartments in macrophages. BMC Biol. 11, 89 (2013).

31. Rodrigues, V., Ruffin, N., San-Roman, M. & Benaroch, P. Myeloid Cell Interaction with HIV: A Complex Relationship. Front Immunol 8, 1698 (2017).

32. Orenstein, J. M., Meltzer, M. S., Phipps, T. & Gendelman, H. E. Cytoplasmic assembly and accumulation of human immunodeficiency virus types 1 and 2 in recombinant human colony-stimulating factor-1-treated human monocytes: an ultrastructural study. J Virol 62, 2578–2586 (1988).

33. Raposo, G. et al. Human macrophages accumulate HIV-1 particles in MHC II compartments. Traffic 3, 718–729 (2002).

34. Deneka, M., Pelchen-Matthews, A., Byland, R., Ruiz-Mateos, E. & Marsh, M. In macrophages, HIV-1 assembles into an intracellular plasma membrane domain containing the tetraspanins CD81, CD9, and CD53. J Cell Biol 177, 329–341 (2007).

35. Jouve, M., Sol-Foulon, N., Watson, S., Schwartz, O. & Benaroch, P. HIV-1 buds and accumulates in ‘nonacidic’ endosomes of macrophages. Cell Host & Microbe 2, 85–95 (2007).

36. Bèrre, S. et al. CD36-specific antibodies block release of HIV-1 from infected primary macrophages and its transmission to T cells. Journal of Experimental Medicine 210, 2523–2538 (2013).

37. Freed, E. O. HIV-1 assembly, release and maturation. Nat Rev Micro 13, 484–496 (2015).

38. Jones, D. M. & Padilla-Parra, S. Imaging real-time HIV-1 virion fusion with FRET-based biosensors. Nature Publishing Group 5, 13449 (2015).

39. Miyauchi, K., Kim, Y., Latinovic, O., Morozov, V. & Melikyan, G. B. HIV enters cells via endocytosis and dynamin-dependent fusion with endosomes. Cell 137, 433–444 (2009).

40. Hammonds, J. E. et al. Siglec-1 initiates formation of the virus-containing compartment and enhances macrophage-to-T cell transmission of HIV-1. PLoS Pathog 13, e1006181 (2017).

41. Chertova, E. et al. Proteomic and biochemical analysis of purified human immunodeficiency virus type 1 produced from infected monocyte-derived macrophages. J Virol 80, 9039–9052 (2006).

42. Ganor, Y. HIV-1 reservoirs in urethral macrophages of patients under suppressive antiretroviral therapy. Nature Microbiology1–48 (2019).

43. Akiyama, H., Ramirez, N.-G. P., Gudheti, M. V. & Gummuluru, S. CD169-mediated trafficking of HIV to plasma membrane invaginations in dendritic cells attenuates efficacy of anti-gp120 broadly neutralizing antibodies. PLoS Pathog 11, e1004751 (2015).

44. Honke, N. et al. Enforced viral replication activates adaptive immunity and is essential for the control of a cytopathic virus. Nat Immunol 13, 51–57 (2011).

45. Oswald-Richter, K. et al. HIV infection of naturally occurring and genetically reprogrammed human regulatory T-cells. PLoS Biol 2, E198 (2004).

46. Oswald-Richter, K., Grill, S. M., Leelawong, M. & Tseng, M. Identification of a CCR5-expressing T cell subset that is resistant to R5-tropic HIV infection. PLoS … 3, e58 (2007).

47. Manel, N. et al. A cryptic sensor for HIV-1 activates antiviral innate immunity in dendritic cells. Nature 467, 214–217 (2010).

48. Cavrois, M., de Noronha, C. & Greene, W. C. A sensitive and specific enzyme-based assay detecting HIV-1 virion fusion in primary T lymphocytes. Nat Biotech 20, 1151–1154 (2002).

49. Manel, N. et al. A cryptic sensor for HIV-1 activates antiviral innate immunity in dendritic cells. Nature 467, 214–217 (2010).

