## Supplemental Information for "Constitutive Siglec-1 expression by human dendritic cell precursors enables HIV-1 replication and transmission"

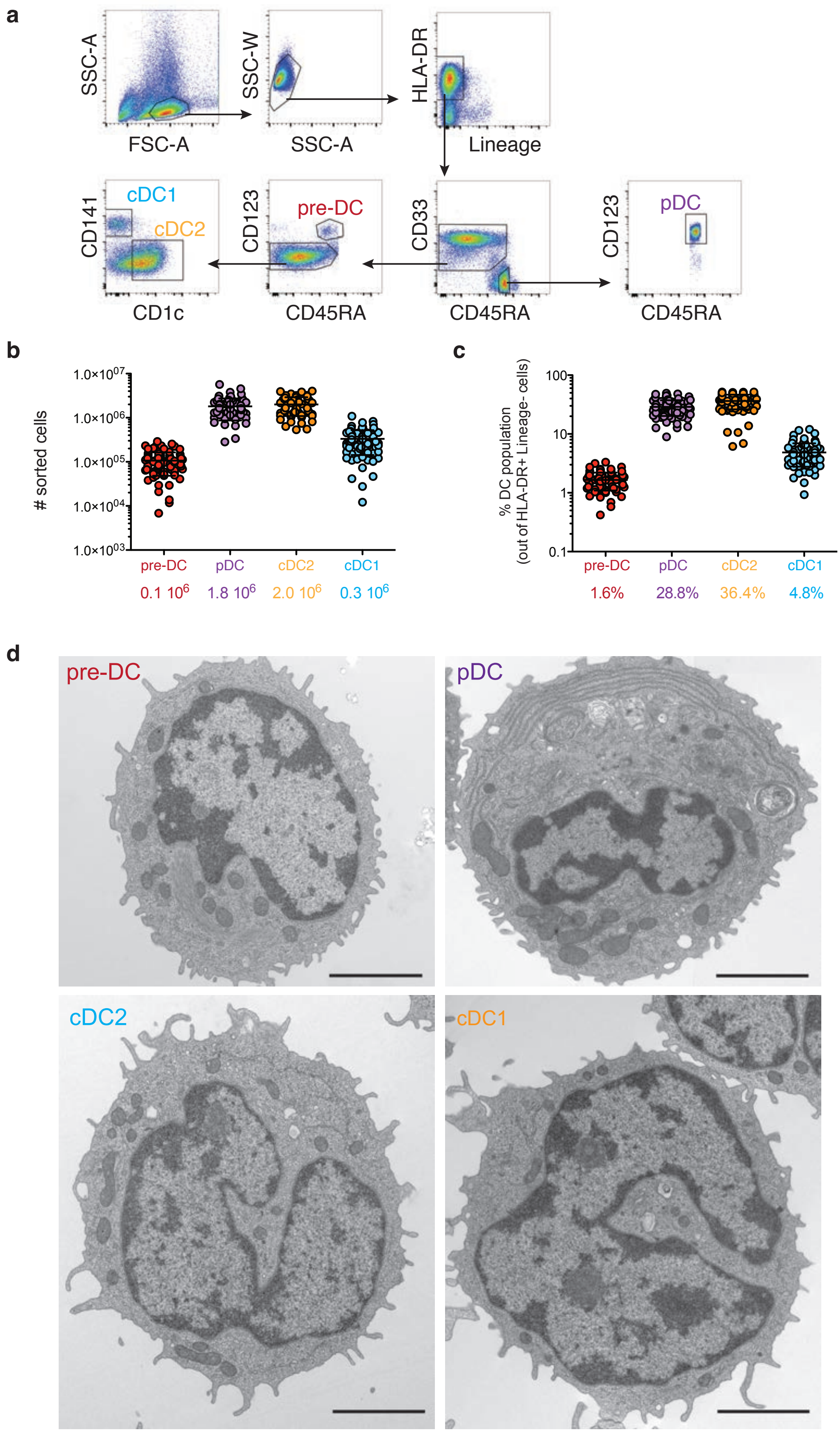


**Figure S1.** Related to Figure 1. Characterization of the sorted blood DC populations.

**a,** Gating strategy used to purify the four populations of DC from the blood after magnetic bead-enrichment of pan DC from PBMC.
**b,** Absolute numbers of cells obtained after sorting for each of the four DC subsets (n= 84 donors). Data represent individual donors with mean ± S.D.
**c,** Percentages of the different subsets among total HLA-$DR^{+}Lin^{-}$ DC (n=84 donors). Data represent individual donors with mean ± S.D.
**d,** Electron microscopy analysis of the four DC subsets freshly sorted from blood. Representative epon sections are presented. Bar scale: 2 μM.

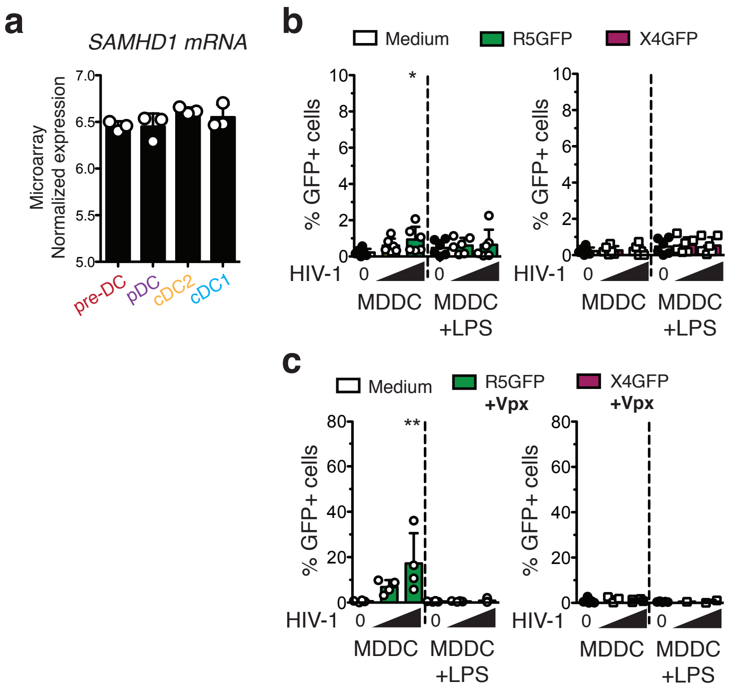


**Figure S2.** Related to Figure 1. Susceptibility of MDDC to HIV-1 infection.

**a,** mRNA levels of SAMHD1 in the freshly sorted DC subsets, measured by RNA seq, (n=3 independent donors combined in 1 experiment). Individual donors are displayed with bars representing mean ± S.D.

**b,** Quantification of GFP expression in MDDC and LPS-activated MDDC infected for 48 h with HIV-1 R5GFP or X4GFP alone or **c,** supplemented with Vpx, n=4 from 4 independent experiments.

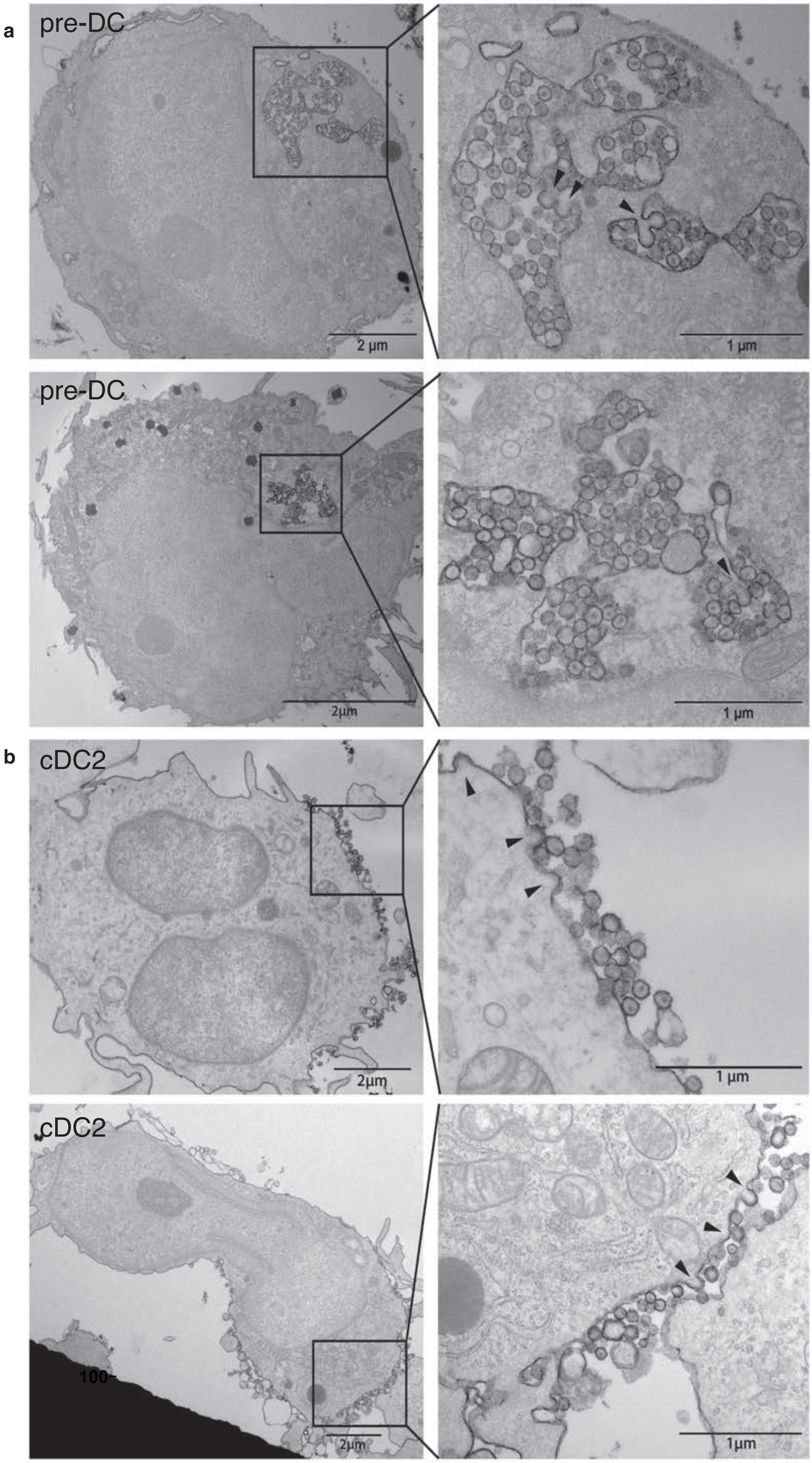

a
pre-DC
2 μm
1 μm
pre-DC
2μm
1 μm
b
cDC2
2μm
1 μm
cDC2
100-
2μm
1μm

**Figure S3.** Related to Figure 4. Ultrastructural analysis of HIV-1-infected pre-DC and cDC2.

**a,** EM profiles of sorted pre-DC or **b,** cDC2 that were infected for 48 h with HIV-1 R5+Vpx. Samples were prepared and processed for EM analysis as in Figure 4a and 4b. Immature virions possess a darker periphery and an electron lucent centre while mature virions contain an electron dense zone in their centre[32,33] revealing the presence of the capsid shell with sometimes its typical conical shape depending on the angle of cut in pre-DC VCC.

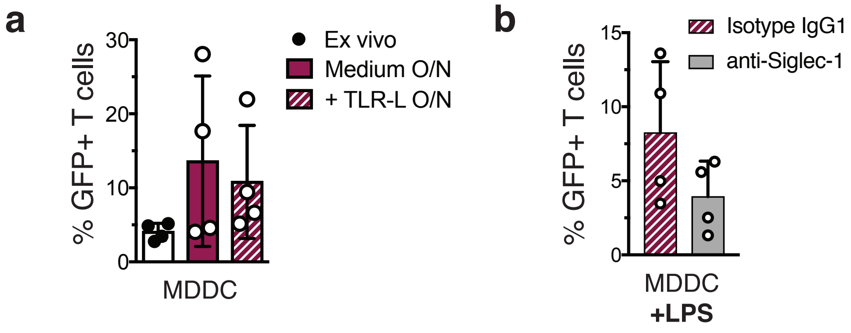


**Figure S4.** Related to Figure 5. Activated MDDC transmit HIV-1 in *trans* to activated CD4+T cells through Siglec-1.

**a,** MDDC were assayed for *trans*-infection as in Figure 5A. GFP expression in activated T cells co-cultured directly with MDDC or following overnight culture in medium or in the presence of LPS. Prior to CD4+T cell co-culture, DC populations were exposed to HIV-1 X4GFP (without Vpx) for 2 h and washed.

**b,** GFP expression in activated T cells co-cultured with LPS-activated MDDC. Prior to CD4+T cell co-culture, MDDC populations were exposed to an anti-Siglec-1 mAb or its isotype control for 30 min and then exposed to HIV-1 X4GFP (without Vpx) for 2 h and washed. For (a) and (b), quantification of $GFP^+$ T cells ($CD3^+$) was performed by flow cytometry after a 48h co-culture, n=4 independent donors. Individual donors are displayed with bars representing mean ± S.D.

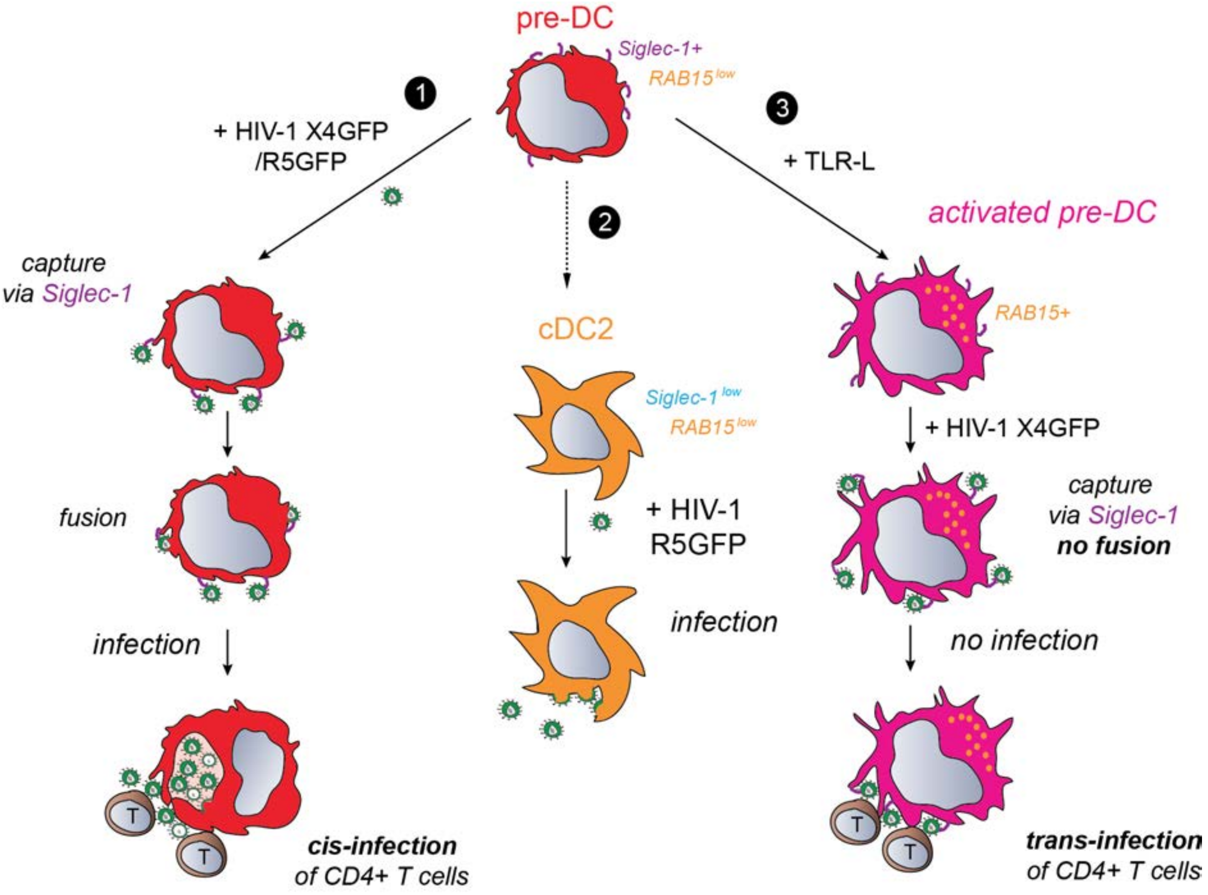


**Figure S5**. Graphical Abstract. pre-DC can contribute to viral spreading via multiple routes.

**1**, pre-DC can capture HIV-1 via their surface Siglec-1 leading to fusion and infection. They produce new viral particles in apparently intracellular virus-containing compartment and can transmit the infection in *cis* to activated T cells. **2**, pre-DC can differentiate into cDC2 that are susceptible to HIV-1 R5-tropic viral infection. HIV-1-infected cDC2 produce viral particles at their plasma membrane. **3**, Activated pre-DC can capture HIV-1 via Siglec-1 and efficiently transfer in *trans* viral particles to activated T cells, which become infected while activated pre-DC are resistant to viral fusion and infection, in association with their increased RAB15 expression.
